# Chronic semaglutide treatment attenuates cue-induced reinstatement of cocaine-seeking and cocaine-induced stereotypy in male and female rats

**DOI:** 10.64898/2026.09.21.753157

**Authors:** Christopher A. Turner, Natalia Morales Pagán, Daniela Pereira, Eileen Y. Sun, Rachel E. Himelhoch, Stephen E. Chang, Shelly B. Flagel

## Abstract

Following cocaine self-administration, male and female rats received daily semaglutide during a 30-day abstinence period before cue- and cocaine-induced reinstatement tests. Semaglutide attenuated cue-induced reinstatement and cocaine-induced stereotypy, although operant responding was elevated following cocaine priming. These findings support further investigation of semaglutide as a treatment for cocaine use disorder.

## Introduction

Glucagon-like peptide-1 (GLP-1) receptor agonists, such as semaglutide, are best known for reducing food intake and sustaining weight loss [1]. These medications also reduce alcohol craving and consumption [2], and emerging evidence suggests they reduce the risk of substance use disorders, including cocaine use disorder [3]. Consistent with these findings, preclinical studies show that acute GLP-1 receptor agonist treatment reduces cocaine-taking and -seeking [4,5]. However, repeated treatment may better model the prolonged exposure associated with clinical use.

A recent study found that repeated semaglutide shifted reward preference in rats from cocaine toward food under concurrent access [6], but whether chronic semaglutide attenuates relapse-relevant behaviors, including cue- and cocaine-induced reinstatement, remains unknown. We therefore assessed the effects of chronic semaglutide treatment, initiated during abstinence after cocaine self-administration, on cue- and cocaine-induced reinstatement in rats.

## Materials & Methods

Subjects were male and female Sprague-Dawley rats weighing 250-325 g upon arrival. All procedures conformed to the *Guide for the Care and Use of Laboratory Animals* (National Research Council, 2011) and were approved by the University of Michigan Institutional Animal Care and Use Committee. Figure 1A shows the timeline; details are in Supplementary Information.

**Figure 1.**
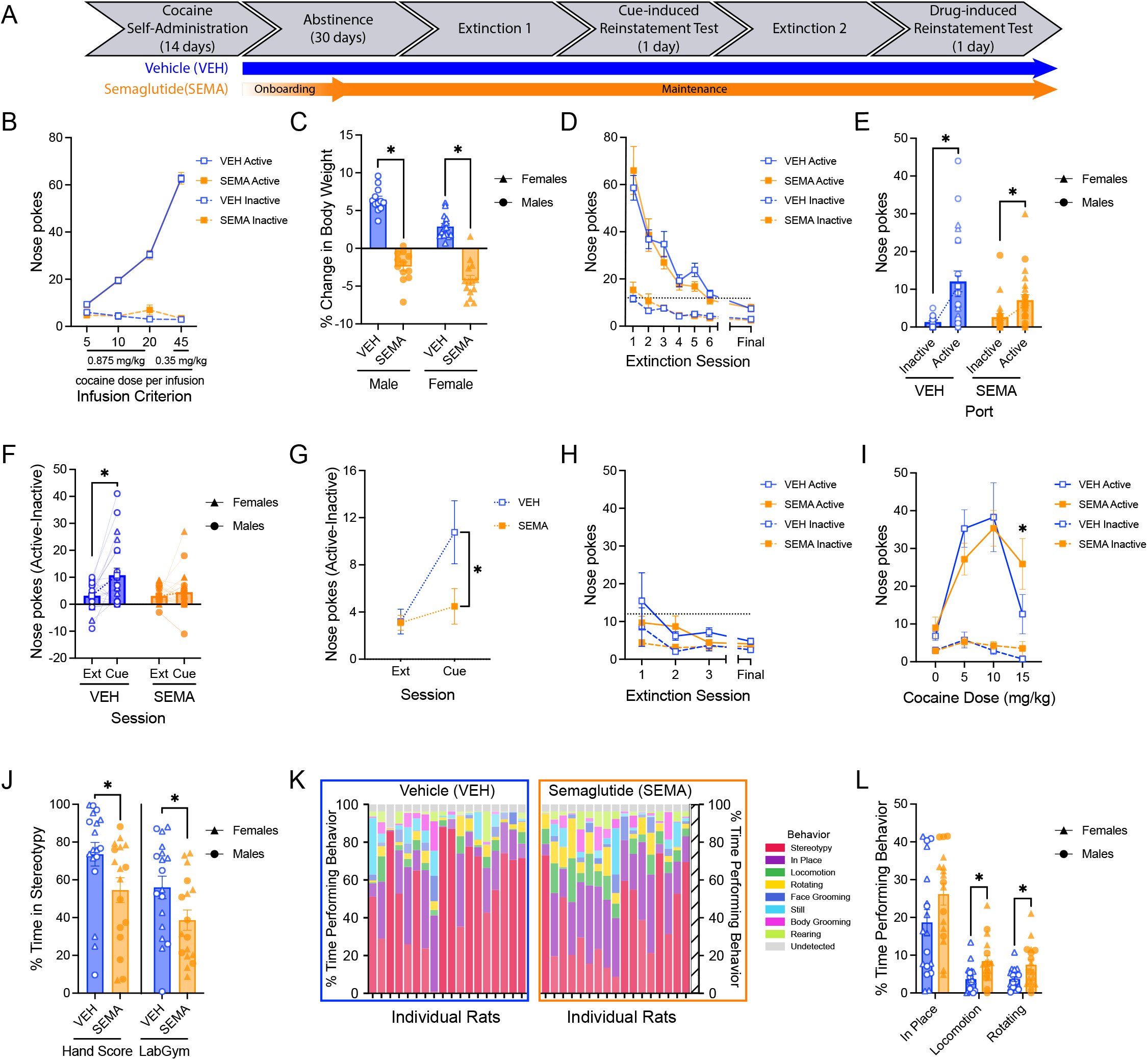
Unless noted, line graphs show group means ± SEM of active-port (solid line) and inactive-port (dashed line) nose pokes, and bars depict group means ± SEM with individual data points shown. Females are indicated by triangles and males by circles. (A) Experimental timeline: vehicle (VEH) or semaglutide (SEMA) treatment began on abstinence day 5 (dose-escalated from 7 µg/kg to a maintenance dose of 70 µg/kg) and continued for the duration of the study. (B) Nose pokes during acquisition of cocaine self-administration across infusion criteria (IC). Rats responded more in the active port as IC increased (*p* < 0.001). (C) Percent change in body weight from baseline after 30-days of SEMA or VEH treatment. SEMA-treated males (^*^*p* < 0.001) and females (^*^*p* < 0.001) showed a greater reduction in body weight compared to their VEH-treated counterparts. (D) Nose pokes across the extinction sessions preceding cue-induced reinstatement. Drug-seeking behavior decreased over time (*p* < 0.001). (E) Nose pokes made during the cue-induced reinstatement test. Dashed lines indicate the active/inactive response ratio. Both groups showed greater responding in the active port than in the inactive port (^*^*p* < 0.001). (F) Active-inactive difference scores for the final extinction session versus the cue-induced reinstatement test. Only VEH-treated animals showed a significant increase in their difference scores during the cue-test relative to extinction (^*^*p* = 0.007). (G) During cue-induced reinstatement, VEH-treated rats had a significantly greater active-inactive difference score than SEMA-treated rats (^*^*p* = 0.049). (H) Nose pokes across the extinction sessions preceding cocaine-induced reinstatement. Overall responding decreased over time (*p* = 0.007). (I) Nose pokes across the cocaine dose-response curve for cocaine-induced reinstatement. At 15 mg/kg, SEMA-treated rats responded significantly more than VEH-treated rats (^*^*p* = 0.026). (J) Percent time in stereotypy following 15 mg/kg cocaine, scored manually (left) and via LabGym (right). Both methods showed that SEMA attenuated stereotypy (HandScore: ^*^*p* = 0.027; LabGym: ^*^*p* = 0.032). (K) Representative LabGym raster plot depicting percentage of time performing each of the eight classified behaviors (stereotypy, in-place, locomotion, rotating, face grooming, still, body grooming, and rearing; below-threshold classifications labeled undetected), averaged across nine 30-s subsamples per animal (columns); color intensity reflects classifier confidence. (L) Percent time spent performing the most frequently detected LabGym-classified behaviors at 15 mg/kg. Treatment groups differed significantly in locomotion (^*^*p* = 0.005) and rotating (^*^*p* = 0.016).

## Cocaine Self-administration

Rats self-administered cocaine through indwelling jugular catheters. An active-port response produced an intravenous cocaine infusion (0.875 mg/kg/infusion, 50 µL over 2.8 s) paired with a 20-s port-light cue, and inactive-port responses were without consequence. To standardize the number of cocaine-cue pairings, infusion criteria (IC) limited infusions across 14 daily sessions: 3 sessions at IC5, 4 at IC10, 3 at IC20, and 4 at IC45. During IC45, cocaine was reduced to 0.35 mg/kg/infusion to elevate operant responding.

## Abstinence and Semaglutide Treatment

Daily subcutaneous vehicle (VEH) or semaglutide (SEMA) injections began on abstinence day 5, and continued throughout abstinence, extinction, and testing, resulting in at least 30 days of treatment before cue-induced reinstatement. SEMA dosing began at 7 µg/kg and increased by 7 µg/kg daily to a maintenance dose of 70 µg/kg [7].

## Extinction

Following 30 days of abstinence, rats returned to the self-administration chambers for 45-min extinction sessions during which responses in either port had no consequence. Training continued until rats made ≤12 active-port responses during each of two consecutive sessions. Rats failing to meet this criterion before reinstatement testing were excluded from corresponding analyses.

## Cue-induced Reinstatement

Cue-induced reinstatement was tested ≈1 h after the last extinction session. During the 45-min test, active-port responses presented the cocaine-associated cue light but no cocaine.

## Cocaine-induced Reinstatement

After additional extinction, cocaine-induced reinstatement was examined 24 h later using a within-session ascending dose-response procedure. Rats received cocaine (0, 5, 10, 15 mg/kg, i.p.) before successive 45-min blocks over ≈4 h. Behavior was video-recorded and scored manually and with LabGym, a behavior-analysis platform [8].

## Statistical Analyses

Statistical details are in the Supplementary Information. Sex differences were observed only in body weight; results for other outcomes are reported across sexes.

## Results

Sample sizes for acquisition, body weight, and extinction were VEH: n=27 (15F/12M) and SEMA: n=26 (14F/12M). After attrition, cue-induced reinstatement sample sizes were VEH: n=21 (10F/11M) and SEMA: n=23 (14F/9M). Cocaine-induced reinstatement was limited to cohorts 2 and 3, with sample sizes of VEH: n=18 (11F/7M) and SEMA: n=17 (10F/7M).

## Cocaine self-administration acquisition

Acquisition of cocaine self-administration was indicated by increased responding across training (effect of IC, F_3,61.445_ = 246.434, *p* < 0.001) and greater active-than inactive-port responding (effect of Port, F_1,134.154_ = 2528.516, *p* < 0.001; IC x Port interaction, F_3, 86.582_ = 660.109, *p* < 0.001; Fig. 1B). Treatment groups were counterbalanced using acquisition measures.

## Chronic semaglutide reduced body weight

Chronic semaglutide reduced body weight in both sexes (effect of Treatment, F_1, 49_ = 223.11, *p* < 0.001; Males: t(22) = 11.805, *p* < 0.001; Females: t(27) = 9.512, *p* < 0.001; Fig. 1C). To determine whether weight differences contributed to reinstatement performance, we correlated body weight with responding. Body weight at the time of testing was not correlated with cue- or cocaine-induced responding (data not shown).

## Chronic semaglutide did not affect the extinction of cocaine-seeking

Semaglutide did not affect extinction before cue-induced (Fig. 1D) or cocaine-induced reinstatement (Fig. 1H), even when the first 5 min of initial extinction were analyzed separately (data not shown).

## Chronic semaglutide attenuated cue-induced reinstatement of cocaine-seeking

During cue-induced reinstatement, both treatment groups responded more in the active than inactive port (effect of Port, Wald χ^2^(1) = 55.14, *p* < 0.001; Fig. 1E). However, VEH-treated rats showed a 9-fold difference between active and inactive responding, whereas SEMA-treated rats exhibited a 2.7-fold difference (Treatment x Port interaction, Wald χ^2^(1) = 7.99, *p* = 0.005; Fig. 1E).

To assess responding above the extinction baseline, active-inactive difference scores were compared between the last extinction session preceding cue-induced reinstatement and the test session (Figs. 1F, 1G). Only VEH-treated rats showed cue-induced reinstatement (Treatment x Session interaction, F_1, 42_ = 4.668, *p* = 0.036; Fig. 1F), as indicated by a significant increase in the active-inactive difference score from extinction to reinstatement in VEH-(t(20) *=* 3.205, p = 0.007) but not SEMA-treated rats (Fig. 1F). In agreement, during reinstatement, the active-inactive response difference was greater in VEH-than SEMA-treated rats (Welch’s t-test: t(31.863) = 2.094, *p* = 0.049; Fig. 1G).

These effects were unlikely to be attributable to nonspecific suppression of activity because inactive-port responding during the cue-induced reinstatement test did not significantly differ between treatment groups (t(42) = 1.363, *p* = 0.180; Fig. 1E), and baseline locomotor activity prior to cocaine-induced reinstatement was comparable between groups (t(33) = 1.354, *p* = 0.185; data not shown).

## Cocaine-induced reinstatement of cocaine-seeking

Rats responded more in the active than inactive port during cocaine-induced reinstatement (effect of Port, F_1, 264_ = 167.812, *p* < 0.001; Fig. 1I). Active-port responding was significantly greater after 5 and 10 mg/kg cocaine than after 0 mg/kg, but decreased at 15 mg/kg, generating an inverted-U dose-response curve (effect of Dose, F_3, 264_ = 27.526, *p* < 0.001; Dose x Port interaction, F_3, 264_ = 11.058, *p* < 0.001; Fig. 1I). Across doses, responding differed between VEH- and SEMA-treated rats (Treatment x Dose interaction, F_3, 264_ = 3.637, *p* = 0.013), regardless of port (Fig. 1I). At 15 mg/kg, SEMA-treated rats responded more than VEH-treated rats (*p* = 0.026; Fig. 1I); however, the Treatment x Port interaction was not significant, indicating increased overall responding rather than selective active-port responding.

## Chronic semaglutide attenuated cocaine-induced stereotypy

Video recordings following the 15 mg/kg dose were scored manually and using LabGym [8]. Manual scoring focused on stereotypy, characterized by repetitive movements such as head swaying, commonly observed following repeated psychostimulant exposure [9]. Both methods showed that chronic semaglutide significantly attenuated the percentage of time spent displaying stereotypy (Handscore: U = 86, *p* = 0.027; LabGym: U = 88, *p* = 0.032; Fig. 1J). Figure 1K shows a plot of LabGym classifications. Relative to VEH-treated rats, SEMA-treated rats spent significantly more time locomoting (t(33) = 2.984, *p* = 0.005) and rotating (Welch’s t-test: t(22.975) = 2.603, *p* = 0.016; Fig. 1L), suggesting that semaglutide shifted the behavioral response to high-dose cocaine from stereotypy toward locomotor activity.

## Discussion

Chronic semaglutide attenuated cue-induced reinstatement of cocaine-seeking and cocaine-induced stereotypy. The incentive-sensitization theory of addiction proposes that repeated drug use induces neural adaptations that sensitize the brain to both the direct effects of the drug (psychomotor sensitization) and drug-associated cues (incentive sensitization), allowing subsequent exposures to trigger pathological drug-seeking [10,11]. Our findings suggest that semaglutide attenuates the expression of incentive and psychomotor sensitization, although further investigation is needed to confirm this interpretation. Additionally, modest cue-induced responding warrants replication under conditions producing more robust reinstatement [12].

SEMA-treated rats showed greater operant responding at the highest cocaine dose during cocaine-induced reinstatement. This likely reflects reduced competition from stereotypy: SEMA-treated rats exhibited greater locomotion and were able to respond, whereas stereotypy interfered with responding in VEH-treated rats. Rather than indicating enhanced cocaine seeking, the increase in responding may reflect a semaglutide-induced shift in the behavioral response to cocaine from stereotypy toward locomotion. Thus, both findings are consistent with attenuated responses to cocaine and cocaine-associated cues. Because responding during reinstatement did not result in cocaine delivery, whether attenuated cocaine effects promote compensatory intake requires further testing.

These findings are consistent with reports that acute GLP-1 receptor agonist treatment attenuates cue- and cocaine-primed reinstatement [4,5]. Notably, attenuated cue-triggered cocaine seeking contrasts with our previously reported enhancement of responding for a food reward and associated cue following chronic semaglutide [13], suggesting reward-specific effects on incentive motivation. Consistent with this interpretation, Heslep et al. found that repeated semaglutide affected cocaine reinforcement more than food reinforcement [6]. These effects may reflect interactions between cocaine-induced adaptations in mesolimbic dopamine signaling [14] and GLP-1 receptor agonist modulation of this system [15], potentially altering psychomotor sensitization to cocaine and the incentive salience of cocaine-associated cues while differentially affecting motivation for food.

In summary, chronic semaglutide attenuated cue-induced reinstatement of cocaine-seeking and cocaine-induced stereotypy, potentially reflecting reduced incentive and psychomotor sensitization, respectively. Semaglutide may offer a novel strategy for reducing relapse vulnerability in cocaine use disorder, warranting further investigation.

## Acknowledgments

The authors thank members of the Flagel Laboratory for their support and insight regarding this research, and Drs. Kent Berridge, Jonathan Morrow, and Terry Robinson for comments on earlier drafts of this manuscript.

## Supplementary Information

### Supplementary Materials and Methods

#### Subjects

After arrival, rats acclimated undisturbed for five days. Rats were housed in a climate-controlled vivarium on a 12-h light:dark cycle with *ad libitum* food and water for the duration of the study. Rats were pair-housed until catheterization surgery, after which they were single-housed to protect implants. Three cohorts of rats were used, with all testing occurring between 11:00 and 15:00 h. Only rats meeting acquisition criteria were included in the analyses.

Subsequent attrition resulted from failure to meet extinction criteria or technical issues. Cocaine-induced reinstatement was conducted only in cohorts 2 and 3.

#### Drugs

Cocaine HCl (Mallinckrodt, St. Louis, MO) dissolved in 0.9% sterile saline was used for self-administration and drug-induced reinstatement. Semaglutide (SEMA; Astatech) was dissolved in 44 mM sodium phosphate dibasic, 70 mM NaCl, and 0.007% Tween 20, which served as the vehicle solution [1].

#### Apparatus

Testing occurred in standard Med Associates operant chambers (Med Associates, St.

Albans, VT, USA; 20.5 × 24.1 cm floor area, 29.2 cm high) enclosed in sound-attenuating boxes, with a ventilation fan providing white noise.

#### Pavlovian Conditioned Approach (PavCA)

Rats underwent PavCA training to phenotype them as sign-trackers, goal-trackers, or intermediate responders, as previously described [2]. These data are not reported because the sample size per phenotype was insufficient.

#### Locomotor Response to Novelty

Following PavCA, locomotor response to novelty was assessed for 60 min in photobeam-equipped chambers [3]; these data are not included in the present study.

#### Surgical Procedures

Following PavCA and locomotor testing, rats received indwelling jugular catheters for self-administration as previously described [4,5]. Catheters were flushed daily with 0.1 mL of heparin (100 U/mL) and gentamicin (1 mg/mL) to maintain patency and prevent infection. Patency was tested before and after the 14-day self-administration paradigm with 0.1 mL of 50 mg/mL methohexital sodium delivered intravenously. Animals losing muscle tone within 3 s were considered patent. Only animals with patent catheters after self-administration were included in the final analyses.

#### Self-administration

The testing chambers were configured for self-administration procedures. Two nose ports were placed on the left and right sides of the chamber, 6 cm above the grid floor. One port was designated the active port and the other the inactive port. The active port was placed on the opposite side of the lever-cue used during PavCA to minimize side bias.

One minute after the start of the session, the house light turned on, and the active-port cue light was illuminated for 20 s. A fixed-ratio-1 (FR1) schedule of reinforcement was used, such that one entry into the active nose port was necessary for cocaine delivery. After a response, the cue light remained lit for a 20-s time-out period, during which additional responses in the active port were recorded without consequence. All responses in the inactive port were recorded without consequence. Sessions concluded after 3 h or once rats met the infusion criterion (IC) [6]. Rats advanced to the next IC after meeting each criterion for at least two consecutive days, and all rats met each criterion in 3-4 sessions.

#### Semaglutide Treatment

Following self-administration, rats underwent 30 days of forced abstinence. Rats were assigned to vehicle (VEH) or semaglutide (SEMA) groups, counterbalanced for sex, body weight, and self-administration performance (e.g., interinfusion interval, session length), and began daily injections on abstinence day 5. Injections were initially administered in the colony room between 12:00 and 13:00 h. Beginning five days before extinction, administration was delayed by 30 min daily until injections occurred between 15:00 and 16:00 h, after extinction or reinstatement sessions each day. Treatment occurred for at least 30 days before cue-induced reinstatement and continued through subsequent extinction and cocaine-induced reinstatement testing.

#### Extinction Training and Excluded Data

Rats underwent two 45-min extinction sessions per day until meeting extinction criteria (≤12 active-port responses during each of two consecutive sessions). The first extinction period, preceding cue-induced reinstatement, consisted of 8–19 sessions, and the second, preceding cocaine-induced reinstatement, consisted of 4–7 sessions. For cue-induced reinstatement, the final extinction session shown in Figures 1D and 1F occurred ≈60 min prior to the cue test. For cocaine-induced reinstatement, the final extinction session occurred ≈24 h prior to testing. Nine rats failed to meet criteria before cue-induced reinstatement: VEH: n = 6 (5F/1M), SEMA: n = 3 (3M). Two rats failed to meet extinction criteria prior to cocaine-induced reinstatement: VEH: n = 2 (1F/1M).

#### Video Scoring

##### Hand Scoring

Videos of the 15 mg/kg cocaine-primed reinstatement session were manually scored for stereotypy. Nine 30-s periods were scored at 5-min intervals from 0 to 40 min, for a total of 4.5 min per subject. Using a stopwatch, experimenters blinded to treatment timed episodes of the same repetitive movement at a fixed location for >2 s, and the timer was stopped when the animal changed location or behavior. The outcome measure was the percentage of observation time spent in stereotypy [(time spent in stereotypy (s)/30 s)^*^100]

##### LabGym Video Analysis

A categorizer was trained in LabGym [7] to classify the following behaviors using animation and pattern images generated from 35 videos (30 fps)

*Stereotypy* – repetitive, spatially restricted movements, predominantly of the head and forepaws (sniffing, gnawing, licking, head-bobbing), directed at a discrete external location (e.g., a wall or a corner). This differs from self-directed grooming and broader in-place ambulatory activity.

*In-place* – small postural or weight shifts, with no significant translational movement.

*Locomotion* – forward movement involving all four limbs with minimal turning, producing a clear forward displacement trail.

*Rotating* – a full rotation returning the animal to or near its original position; partial rotations were rejected.

*Face grooming* – repetitive paw-to-face/head contact.

*Still* – absence of movement outside that caused by respiration.

*Body grooming* – bending or twisting to reach/lick the flank and hindlimb scratching.

*Rearing* – hindlimb support with forepaws lifted and a clear vertical posture; stand-up/drop-down transitions were accepted; brief paw lifts were rejected.

Frames labeled undetected fell below the assigned confidence threshold (Fig. 1K).

#### Statistical Analyses

Sex was initially included in each model but removed when no main effect of sex or interaction involving sex was detected; therefore, behavioral results are presented across sexes.

Acquisition and extinction data were analyzed using linear mixed models (LMMs), with factors of Treatment (VEH vs. SEMA), Time (ACQ: IC; EXT: Session), and Port (Active vs. Inactive). The covariance structure was selected using the lowest Akaike’s information criterion for each dataset. Average percent change in body weight from baseline was analyzed using a two-way ANOVA with factors of Treatment (VEH vs. SEMA) and Sex (Female vs. Male). Cue reinstatement and cocaine reinstatement at 15 mg/kg were analyzed using generalized estimating equations (GEE), with factors of Port (Active vs. Inactive) and Treatment (VEH vs. SEMA). Cue reinstatement vs. final extinction data (active-inactive nose pokes) were analyzed using a repeated-measures two-way ANOVA, with factors of Treatment (VEH vs. SEMA) and Session (Ext vs. Cue). Cocaine reinstatement dose-response data were analyzed using a generalized linear mixed model (GLMM), with factors of Treatment (VEH vs. SEMA), Dose (0, 5, 10, 15 mg/kg), and Port (Active vs. Inactive). Manual and LabGym stereotypy data were analyzed independently using unpaired Mann-Whitney tests. Any statistically significant main effects and interactions were followed up with *post hoc* comparisons (t-tests) with Bonferroni corrections as appropriate. A Welch’s t-test was used when data violated the assumption of homogeneity of variances.

